# rPinecone: Define sub-lineages of a clonal expansion via a phylogenetic tree

**DOI:** 10.1101/404624

**Authors:** Alexander M. Wailan, Francesc Coll, Eva Heinz, Gerry Tonkin-Hill, Jukka Corander, Nicholas A. Feasey, Nicholas R. Thomson

## Abstract

The ability to distinguish between pathogens is a fundamental requirement to understand the epidemiology of infectious diseases. Phylogenetic analysis of genomic data can provide a powerful platform to identify lineages within bacterial populations, and thus inform outbreak investigation and transmission dynamics. However, resolving differences between pathogens associated with low variant (LV) populations carrying low median pairwise single nucleotide variant (SNV) distances, remains a major challenge. Here we present rPinecone, an R package designed to define sub-lineages within closely related LV populations. rPinecone uses a root-to-tip directional approach to define sub-lineages within a phylogenetic tree according to SNV distance from the ancestral node. The utility of this program was demonstrated using genomic data of two LV populations: a hospital outbreak of methicillin-resistant *Staphylococcus aureus* and endemic *Salmonella* Typhi from rural Cambodia. rPinecone identified the transmission branches of the hospital outbreak and geographically-confined lineages in Cambodia. Sub-lineages identified by rPinecone in both analyses were phylogenetically robust. It is anticipated that rPinecone can be used to discriminate between lineages of bacteria from LV populations where other methods fail, enabling a deeper understanding of infectious disease epidemiology for public health purposes.

**DATA SUMMARY:**

1. Source code for rPinecone is available on GitHub under the open source licence GNU GPL 3; (url: https://github.com/alexwailan/rpinecone).
2. Newick format files for both phylogenetic trees have been deposited in Figshare; (url: https://doi.org/10.6084/m9.figshare.7022558)
3. Geographical analysis of the *S*. Typhi Dataset using Microreact is available at https://microreact.org/project/r1IqkrN1X.
4. Accession numbers, meta data and sample lineage results of both datasets used in this paper are listed in the supplementary tables.

**I/We confirm all supporting data, code and protocols have been provided within the article or through supplementary data files. ⊠**

**IMPACT STATEMENT:** Whole genome sequence data from bacterial pathogens is increasingly used in the epidemiological investigation of infectious disease, both in outbreak and endemic situations. However, distinguishing bacterial species which are both very similar and which are likely to come from a small geographical and temporal range presents a major technical challenge for epidemiologists. *rPinecone* was designed to address this challenge and utilises phylogenetic data to define lineages within bacterial populations that have limited variation. This approach is therefore of great interest to epidemiologists as it adds a further level of clarity above and beyond that which is offered by existing approaches which have not been designed to consider bacterial isolates containing variation that only transiently exist, but which is epidemiologically informative. *rPinecone* has the flexibility to be applied to multiple pathogens and has direct application for investigations of clinical outbreaks and endemic disease to understand transmission dynamics or geographical hotspots of disease.

## INTRODUCTION

Advances in Whole Genome Sequencing (WGS) permit the study of bacteria at high resolution and this has created the opportunity to use WGS data to reliably discriminate between bacteria that were previously indistinguishable by alternative means, or to “type” them. Using WGS data to understand the epidemiology of infections presents distinct challenges to those faced by evolutionary biologists, more so when aiming to distinguish lineages of bacterial populations that are only transiently extant.

Single nucleotide variants (SNVs) which arise stochastically may offer no evolutionary benefit to a given bacterial isolate, and therefore may only transiently be present in a population. Despite their lack of evolutionary significance, however, they may be epidemiologically informative in the context of localised outbreaks, especially amongst organisms that show low overall population diversity. Traditional genetic clustering algorithms such as Structure(1) and BAPS(2, 3) are not suitable for sub-typing of low-variant (LV) bacterial populations over small timescales such as less than three years. They assume independence between loci and require 10-100’s of SNV that are shared between different lineages to confidently cluster populations. Novel bioinformatic approaches to identify sub-lineages within LV bacterial populations are therefore needed for epidemiological investigations, especially when technological advancements have made genomic studies of outbreaks increasingly common.

For genomic data, typing tools are used to identify “clusters” or “partitions” to define a group of isolates as a “lineage” or sub-lineage within a population. These three terms are used synonymously in this context. However, to provide clarity we will use the term “sub-lineage” to refer to a defined group of isolates within a population. These typing tools are non-tree based and fall under two broad approaches, distance-based methods which calculate a pairwise distance matrix often using the SNV distances between isolates, and model-based methods which rely on calculating a population genetic model using either Bayesian or maximum likelihood based methods(1). For distance methods samples are considered to be within the same cluster if their SNV distance is less or equal to a specified SNV threshold. An alternative method is the widely used program hierBAPS(3), which when given a multiple sequence alignment attempts to identify the lineages of the sequences that maximises the posterior probability of the hierBAPS model. The model is based on a Multinomial-Dirichlet distribution and assumes independence between SNV sites. The algorithm is applied first to identify an initial set of lineages, which is then iteratively repeated to generate subsequent sub-lineages within each of the initially defined lineages. While both are suitable for populations where long-term historical evolution is present, neither are appropriate options for short-term clonal expansions such as populations emerging from point source outbreaks or locally endemic disease. In the case of Bayesian approaches, the resulting sub-lineages are not able to reflect the phylogenetic data, whereas methods that use a pairwise SNV distance approach are unable to infer direction of evolution or common ancestry between sub-lineages, which is necessary for source and transmission investigations.

Here we present rPinecone, an R package which evaluates a phylogeny using a root-to-tip approach to define sub-lineages according to SNV distances from ancestral nodes. rPinecone also determines if two or more sub-lineages form part of a larger major lineage if they are related by enough ancestral nodes. To demonstrate this approach, rPinecone was tested on a local endemic dataset of *Salmonella enterica* subsp. *enterica* serovar Typhi (*S*. Typhi) from the H58 haplotype(4) as well as a hospital outbreak of methicillin-resistant *Staphylococcus aureus* (MRSA) ST2371(5).

## METHODS

### Phylogenetic tree construction of two case studies

Data from two studies were reanalysed to construct phylogenetic trees for input into rPinecone. Mapping of reads was performed using SMALT v0.7.4 (sanger.ac.uk/science/tools/smalt-0/) and SNVs were identified by using samtools mpileup(6). SNV alignments were generated using SNP-sites(7) then used to generate phylogenetic maximum-likelihood (ML) trees with RAxML(8). In the MRSA case study, the alignment and subsequent rooted ML tree was generated by using the *S. aureus* ST22 isolate HO 5096 0412 (accession number HE681097) as a mapping reference and as an outgroup to root the tree. In the case of the locally endemic *S.* Typhi in Cambodia, *S.* Typhi Genotyphi 4.3.1 isolate 2010_7898 (accession number GCA_001360555) was used as a mapping reference. *S.* Typhi Genotyphi 4.1.1. isolate Mal1017142 (accession number ERR279139)(9) was used as an outgroup to root the ML tree. Methodological details for construction of phylogenetic trees can be found in the supplementary material.

### Ancestral reconstruction of phylogenetic trees

The SNV alignment and ML tree from the reanalysis of both datasets was used as an input for the ancestral reconstruction tool “pyjar” (github.com/simonrharris/pyjar)(10) generating a ML joint ancestral reconstruction (JAR) tree. Each JAR tree was used as an input for rPinecone to define the sub-lineages within respective populations.

### hierBAPS as a comparative benchmark

The R package for hierBAPS (rhierBAPS, available at github.com/gtonkinhill/rhierbaps) was used as a comparative benchmark for rPinecone when analysing the *S*. Typhi dataset, due to its frequent use in bacterial phylogenetics for lineage identification. Input data for this package was the SNV alignment used to generate the phylogenetic ML tree. A maximum level depth of five and maximum cluster number of 30 was used for the analysis.

## RESULTS

rPinecone is a package developed in R and available under the open source licence GNU GPL3 at github.com/alexwailan/rpinecone. Input for rPinecone requires two user-specified integer thresholds for SNV distance and relatability (discussed below), and a rooted phylogenetic tree (Newick format) where the branch lengths are integers corresponding to SNV distance. One type of tree that fits this criterion is a maximum likelihood (ML) joint ancestral reconstruction (JAR) tree.

The rPinecone package has a primary wrapper function to perform the analysis which is performed in two steps: sub-lineage definition and major-lineage definition (Fig. 1).

**Figure 1.**
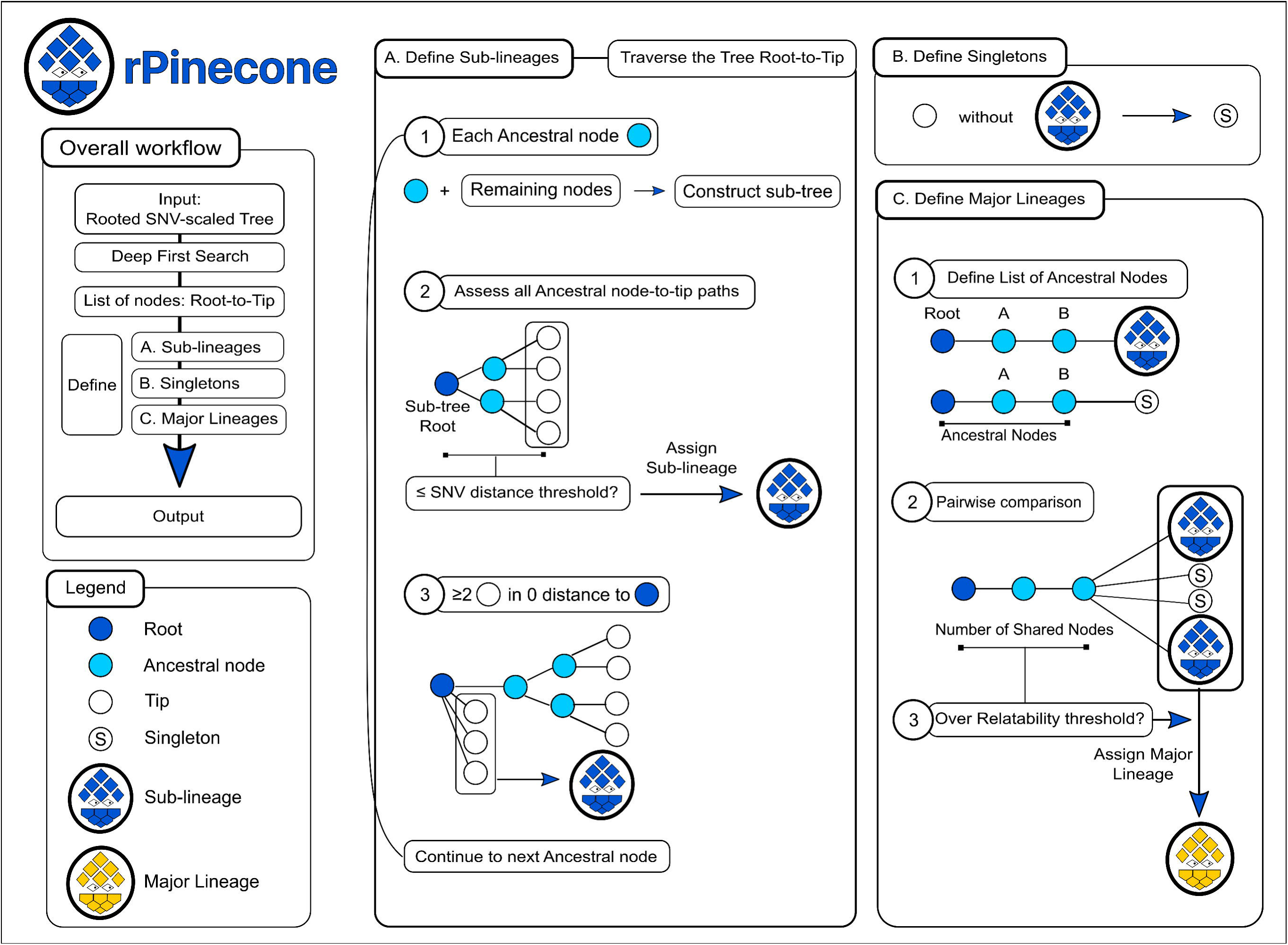
The rPinecone package evaluates a phylogenetic tree in a root-to-tip approach and defines sub-lineages according to single nucleotide variants (SNV) distances from ancestral nodes. (A) Define Sub-lineages. The rPinecone algorithm traverses the tree from root-to-tip in the order of nodes listed by the depth-first search. At each ancestral node, a sub-tree is constructed containing the ancestral and remaining nodes. The ancestral node becomes the root of the sub-tree and all root-to-tip paths are assessed to identify the maximum root-to-tip branch distance. If this distance is below or equal to the user specified SNV distance threshold, the tips of the sub-tree are assigned to a sub-lineage number. If the distance exceeds the threshold and two or more tips are in zero distance to the root of the sub-tree, only the tips in zero distance will be assigned a sub-lineage number. The process is repeated throughout the entire tree. (B) Define Singletons. Tips which have not been assigned a sub-lineage number are assigned a singleton number. (C) Define Major lineages. Using the entire tree, the list of ancestral nodes from the root to each sub-lineage and singleton is compiled. A pairwise comparison of these lists is performed. Sub-lineages and singletons which have the same number of ancestral nodes over the user specified relatability threshold will be defined as part of a major lineage. The root, ancestral nodes, and tips of the tree are denoted by navy, cyan and white circles respectively. Singletons, sub-lineages, major lineages are represented by a ‘S’ labelled circle, a blue pinecone, a gold pinecone.

rPinecone will initially prepare for sub-lineage definition by collapsing zero SNV dichotomies of the tree into polychotomies using the *ape* package(11) for R (v4.1) and performing a depth-first search (DFS)(12) across the resulting tree, to list the tips and ancestral nodes from root-to-tip. Edges are then drawn between nodes where the edge distance is the branch length i.e. the number of SNVs isolates have from an ancestral node on a JAR tree. Using the DPS and SNV branch distance information, rPinecone will then define sub-lineages by traversing the tree in a root-to-tip direction to assess each ancestral node. At each ancestral node, a sub-tree is built with the remaining nodes where the ancestral node becomes the root of the sub-tree. rPinecone uses two methods to define sub-lineages, 1) all root-to-tip paths of the sub-tree are assessed and if the maximum root-to-tip branch distance is equal or below the specified SNV threshold, the remaining tips will be assigned a sub-lineage number and 2) If the maximum distance is over the SNV threshold and there are at least two tips with zero distance from the ancestral node, rPinecone assigns these tips in zero distance a sub-lineage number. The process is repeated throughout the entire tree. Tips that are not assigned a sub-lineage number will be assigned with a singleton number.

After sub-lineages have been declared, their relatability is analysed to determine if they form a larger major lineage. The user specified relatability threshold is an integer used to compare the number of ancestral nodes from the root of the entire tree to each sub-lineage and singleton. Lists of ancestral nodes for each sub-lineage and singleton are compiled. These lists are compared in a pairwise fashion. Sub-lineages and singletons that have the same intersecting ancestral nodes over the relatability threshold are declared to have formed a major lineage within the population, where a major lineage is composed of at least two sub-lineages.

Once sub-lineages and major lineages are defined, rPinecone outputs a list of six variables each able to be used for downstream processes, including; the tree used for analysis, number of sub-lineages, major lineages, singletons identified and number of isolates within the tree. The final variable is a three-column table listing each “Taxa” label of the tree and their respective sub-lineage and major lineage. Furthermore, phylogenetic trees can be displayed using the online display tool iTOL(13). The rPinecone package includes three functions taking the output from the primary rPinecone function. These functions parse the rPinecone output to a file in a format to display in iTOL. Three files can be generated, a “LABELS” file to change the tip labels according to their sub-lineage or singleton number as well as two “DATASET_COLORSTRIP” format files to display the sub-lineages and major lineages as a colour strip/ block (Example displayed in the Fig. 2). To show the utility of rPinecone we perform two case studies on a MRSA hospital outbreak and local endemic dataset of *S.* Typhi.

**Figure 2.**
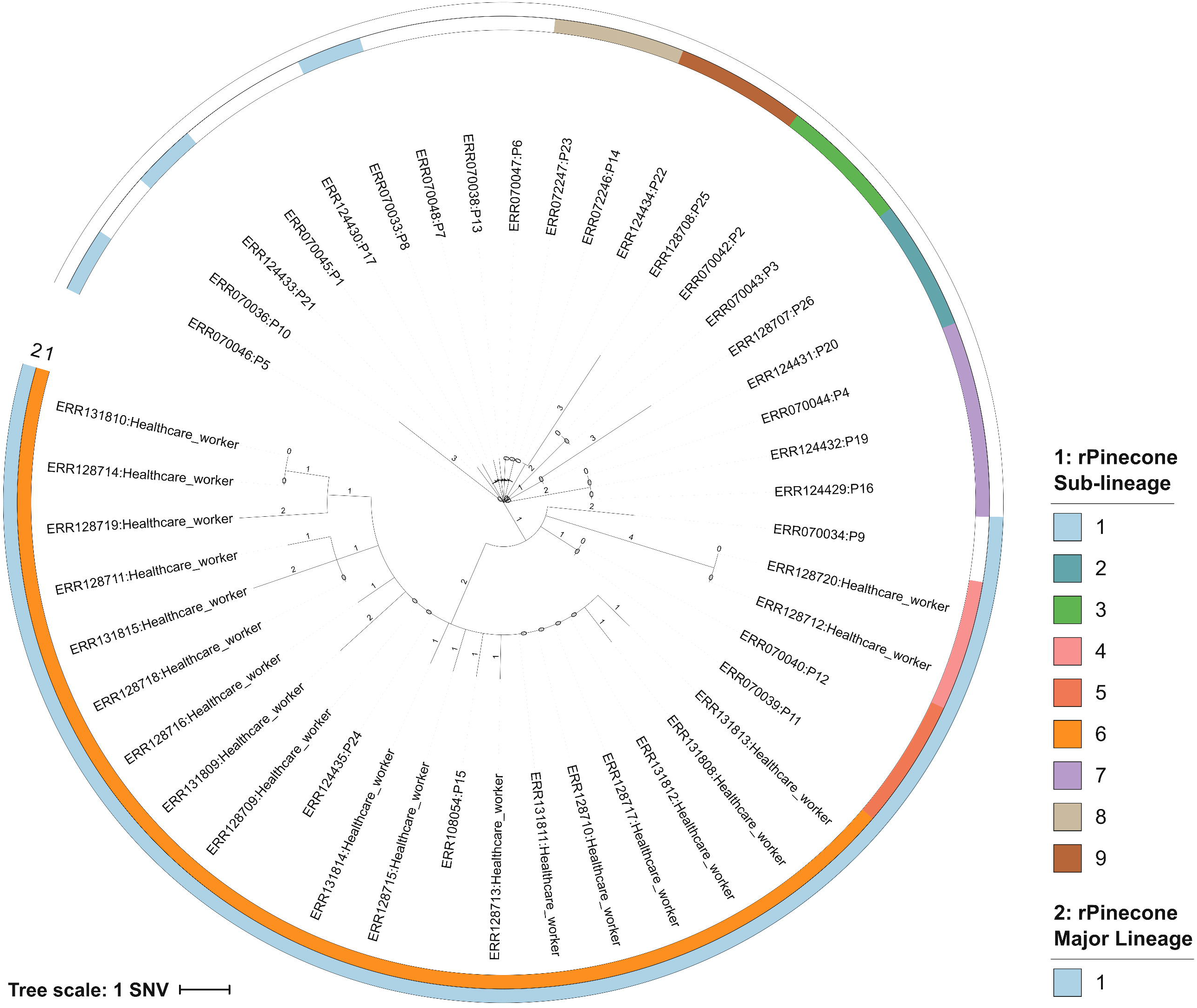
Rooted accelerated transformation tree based on whole genome sequencing of 45 MRSA ST2371 isolates. Outgroup rooted using *S. aureus* ST22 isolate HO 5096. Tips are labelled by the sample’s ERR number followed by the origin of sample, patient (P) or Healthcare worker. Branch lengths are single nucleotide variants scaled. The inner and outer coloured rings correspond to the sub-lineages and, major lineage identified by rPinecone.

**Figure 3.**
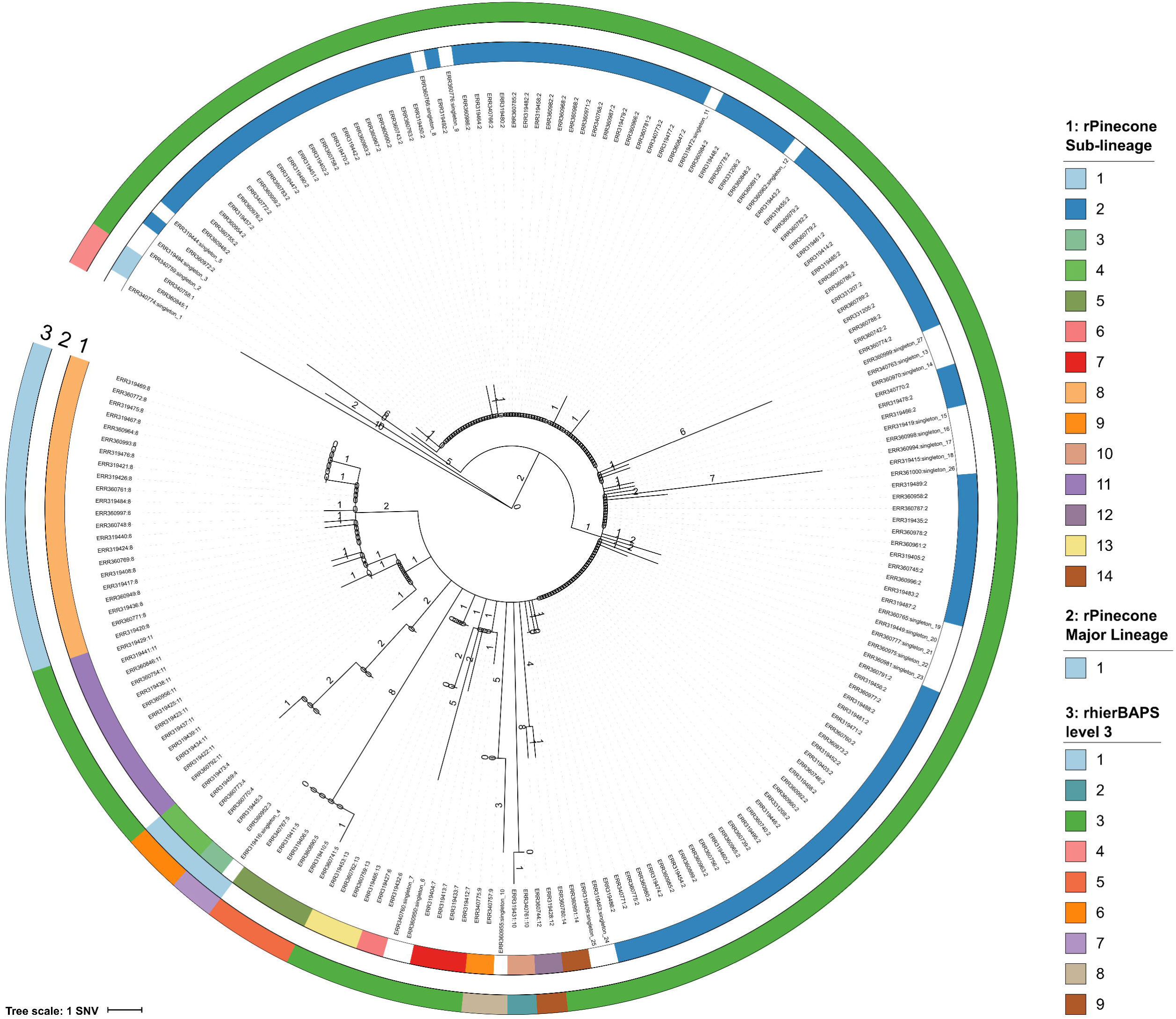
Rooted joint ancestral reconstruction tree based on whole genome sequencing of 203 *S*. Typhi H58 isolates. Outgroup rooted using *S.* Typhi Genotyphi 4.1.1 isolate Mal1017142. Tips are labelled by the sample’s ERR number and rPinecone sub-lineage. Branch lengths are single nucleotide variants scaled. Colour rings one and two correspond to the sub-lineages and major lineage defined by rPinecone. The colour ring three corresponds to clusters defined by rhierBAPs level three.

## HOSPITAL OUTBREAK OF MRSA ST2371 – HEALTHCARE WORKERS WERE COLONISED BY A MAJOR LINEAGE

In 2011 over a 6-month period, the National Health Service Foundation Trust in Cambridge, UK, investigated transmission of MRSA within the hospital neonatal unit and community and described a complex transmission network(5). We reanalysed the data of this study and generated a phylogenetic ML tree. From this analysis the population had a median pairwise distance of five SNVs. The phylogenetic tree (Fig. 2) of the outbreak had a structure that begins with a root isolate (Patient 5), then expands outwards in a “star-burst”-like fashion where each branch of the tree represents a transmission pathway of MRSA between infants and their mothers, other mothers on the ward, and to partners of affected mothers as described in the original study(5). rPinecone identified nine sub-lineages and a major lineage composed of three sub-lineages (25 isolates) using SNV threshold of four and a relatability threshold of three. These sub-lineages corresponded to the transmission pathways determined in the original investigation using epidemiological data. The major lineage represents the sub-lineages of MRSA (rPinecone sub-lineages 4, 5 and 6) carried by colonised healthcare workers which were associated with subsequent episodes of MRSA 64 days after the last MRSA-positive patient left the unit.

The major lineages identified by rPinecone highlight sub-lineages related by ancestral nodes and potentially lineages that have separated from the remaining clonal population. This type of analysis was not performed during the original analysis, however we successfully identified MRSA transmission pathways using rPinecone and provided an additional layer of sub-lineage data to complement the epidemiological data when associating sub-lineages with likely transmission events.

## LOCALLY ENDEMIC *S.* TYPHI IN RURAL CAMBODIA – HIGHER RESOLUTION TO IDENTIFY CIRCULATING SUB-LINEAGES

In the second example, rPinecone distinguished lineages of an *S*. Typhi population circulating in a defined geographical area causing locally endemic disease, as opposed to a clonal outbreak. LV bacterial pathogens responsible for localised, endemic disease present a different challenge; in this context, it is highly likely that multiple, genetically similar sub-lineages will be in circulation, and in order to be epidemiologically informative, phylogenetic analysis must be able to distinguish these lineages despite the low variation between them. Whilst hierBAPS is unable to utilise the low variant SNV information to define lineages, rPinecone can be used in such situations. We demonstrate this by defining sub-lineages in a LV population of *S.* Typhi H58 responsible for endemic typhoid fever in a rural part of Cambodia(4). Reanalysis of these data determined the population to have a median pairwise SNV distance of two. The subsequent phylogenetic tree of the reanalysis can best be described from root-to-tip (Fig. 3). At the root of the tree resides a “root group” of isolates, then a main branch composing of identical isolates which can be referred to as the “primary group” of the clonal expansion, followed by “diverging groups” of isolates and singletons which have diverged from this main branch. Using the reanalysed data, rPinecone identified 14 sub-lineages and a major lineage composed of two sub-lineages (seven isolates) of *S.* Typhi, using a SNV threshold of two and a relatability threshold of three (Fig. 3). 27 isolates were also identified as singletons. In comparison, rhierBAPS identified a maximum of nine sub-lineages at level three and offered no further resolution after further analysis (2, 7, 9, 9 and 9 sub-lineages were identified at levels one through to five [Supplementary Fig.1]).

rPinecone sub-lineages were nested within the sub-lineages defined by rhierBAPS. rPinecone also acknowledges singleton isolates which have accumulated their own “private” SNVs. Therefore, singletons that have individual SNVs may be excluded in the definition of sub-lineages. In contrast, rhierBAPS cannot detect the private SNVs of singletons. This is because rhierBAPs seeks to maximise the posterior probability of a sub-lineage assuming independence between SNV sites, and is less suitable for the identification of singletons. In addition, rhierBAPS also identifies isolates situated across the tree to be members of the same sub-lineage. hierBAPS and other similar programs like Structure are not designed to separate population data that would be observed towards the tips of a phylogenetic tree including those observed with a median pairwise SNV of two as observed here. Rather they aim to cluster isolates into groups that are likely to come from similar source populations. When there are few isolates and few SNVs there is little information to estimate the likely distribution of allele frequencies in these source populations.

In this example, rPinecone provided greater resolution to distinguish sub-lineages than across five different levels of rhierBAPS and distinguished additional sub-lineages. The original study noted their sub-lineages to have significant geographical variation. The rPinecone sub-lineages provided a cross-sectional view of the bacterial population, where some sub-lineages were geographically confined to districts within one province and others span multiple provinces (Supplementary Fig. 2).

## CONCLUSION

Investigators must be able to distinguish isolates to identify epidemiologically informative sub-lineages, however this is difficult for bacterial populations associated with both outbreaks and endemic disease over small temporal or geographical distances when the pathogen has a slow mutation rate and appears largely “clonal”. The ability to make this distinction is critical if WGS data is to inform epidemiological investigation of such pathogens, whether in epidemic, or locally endemic disease. When aiming to geographically locate hotspots of disease transmission, it is critical to be able to incorporate associated geospatial data associated with samples.

rPinecone was designed to analyse LV populations and to be applied to any bacterial species with at least a median SNV distance of two within the population. Using the genomic data of a MRSA hospital outbreak and endemic *S.* Typhi, we demonstrate that rPinecone can be used to identify sub-lineages within LV populations that are reflective of the phylogenetic data. Furthermore, when compared to rhierBAPS, rPinecone identified additional sub-lineages. We highly recommend using heirBAPs for initial analysis of phylogenetic data to understand the bacterial population. Once this has been done, rPinecone will define sub-lineages of LV bacterial populations to set the platform for understanding the transmission dynamics or geographical hotspots of disease.

## AUTHOR STATEMENTS

### Funding information

This project and A.M.W (Postdoctoral Fellow) was supported by a grant from the Bill and Melinda Gates Foundation (OPP1128444). The Wellcome Trust Sanger Institute is core funded by Wellcome Trust grant 206194. JC was funded by ERC grant no. 742158. F.C. is funded by a Wellcome Trust Sir Henry Postdoctoral Fellowship (201344/Z/16/Z).

## ABBREVIATIONS

LV: Low-Variant
SNV: Single Nucleotide Variant
WGS: whole genome sequencing
ML: Maximum-Likelihood
JAR: Joint Ancestral Reconstruction

